# Gut-to-brain regulation of *Drosophila* aging through neuropeptide F, insulin and juvenile hormone

**DOI:** 10.1101/2024.06.26.600832

**Authors:** Jiangtian Chen, Marcela Nouzova, Fernando G. Noriega, Marc Tatar

## Abstract

Dietary restriction slows aging in many animals, while in some cases the sensory signals from diet alone are sufficient to retard or accelerate lifespan. The digestive tract is a candidate location to sense nutrients, where neuropeptides secreted by enteroendocrine cells (EEC) produce systemic signals in response to food. Here we measure how *Drosophila* neuropeptide F (NPF) is secreted into adult circulation by enteroendocrine cells and find that specific enteroendocrine cells differentially respond to dietary sugar and yeast. Lifespan is increased when gut NPF is genetically depleted, and this manipulation is sufficient to blunt the longevity benefit conferred by dietary restriction. Depletion of NPF receptors at insulin producing neurons of the brain also increases lifespan, consistent with observations where loss of gut NPF decreases neuronal insulin secretion. The longevity conferred by repressing gut NPF and brain NPF receptors is reversed by treating adults with a juvenile hormone (JH) analog. JH is produced by the adult *corpora allata*, and inhibition of the insulin receptor at this tissue decreases JH titer and extends lifespan, while this longevity is restored to wild type by treating adults with a JH analog. Overall, enteroendocrine cells of the gut modulate Drosophila aging through interorgan communication mediated by a gut- brain-*corpora allata* axis, and insulin produced in the brain impacts lifespan through its control of JH titer. These data suggest that we should consider how human incretins and their analogs, which are used to treat obesity and diabetes, may impact aging.

**Significance Statement:** Neuropeptide F (NPF) produced in the *Drosophila* gut is an insulin-regulatory hormone (incretin) that is secreted into adult circulation in response to feeding and diet. Suppression of gut NPF extends *Drosophila* longevity, as does knockdown of Neuropeptide F receptors at the insulin-producing medial neurosecretory cells in the brain that control the titer of juvenile hormone. Gut hormones and brain insulin regulate lifespan because they control juvenile hormone titer, which itself is the master endocrine regulator of *Drosophila* aging. Gut NPF modulates Drosophila aging through the integration of nutrient sensing, insulin signaling and juvenile hormone. Given the role of incretin-mimetic drugs to treat diabetes and obesity, it may be time to consider how incretin analogs could impact human aging.

## Main Text

## Introduction

Neuropeptides released from gut enteroendocrine cells (EEC) modulate mammalian physiology (1, 2). Secretin from the intestine regulates gastric acid, pancreatic bicarbonate and water balance (3). Ghrelin produced in the stomach induces appetite through its action upon the hypothalamus, while intestinal Peptide YY (PYY) is anorexigenic (4). Pancreatic insulin secretion is amplified by the gut derived incretins glucagon-like peptide 1 (GLP-1) and glucose-dependent insulinotropic peptide (GIP) (5, 6). Importantly, analogs of GLP-1 are now used to treat diabetes and obesity (7–9).

Aside from mammals, enteroendocrine cells are found in the digestive systems of many animals (10–12). The midgut of *Drosophila melanogaster* contains EEC that produce 24 neuropeptides including allatostatins, Dh31, tachykinin, CCHamides, and Neuropeptide F (NPF) (13–16). While many of these neuropeptides are also produced in the brain (17), when derived just from the gut these peptides can directly modulate intestinal stem cell division, renal function, feeding, metabolism and courtship (18–21). Among these, *Drosophila* NPF is an amidated PP-fold neuropeptide that resembles mammalian Neuropeptide Y (NPY) and Peptide YY (PYY) (22, 23). In addition, the *Drosophila* NPF receptor (NPFR) responds to both NPY and PYY when expressed in *Xenopus* oocytes (24). NPF receptors in adult *Drosophila* are found in the ovary (25), fat body (26), *corpora cardiaca* (27), brain (28) and ventral nerve cord (29). Central to our current study, NPF secreted from the midgut is an incretin that induces secretion of insulin from brain median neurosecretory cells (MNC) (26, 27).

Notably, systemic reduction of insulin signaling extends *Drosophila* lifespan (30–32). Here we hypothesize that gut NPF modulates aging by regulating insulin secretion from median neurosecretory cells (MNC), and that the nutrient responsiveness of gut NPF mediates how dietary restriction extends lifespan. We then test whether gut NPF impacts aging because insulin released from median neurosecretory cells stimulate the *corpora allata* to produce juvenile hormone (JH), where JH itself systemically regulates *Drosophila* aging.

We quantified how dietary yeast and sugar differentially influence the level of NPF secreted into the hemolymph by distinct gut enteroendocrine cells. Inhibition of gut NPF from open-type EECs mediated the longevity conferred by dietary restriction, while inhibition of closed-type EEC extended lifespan independent of diet. Gut-derived NPF localized at insulin-producing medial neurosecretory cells within the brain, and lifespan was increased by blocking NPF receptors exclusively in these neurons. This inhibition impaired insulin secreted from the MNC, which limited JH released from the *corpora allata* (CA). Our previous research demonstrated that lifespan conferred by mutation of *Drosophila* insulin receptors was reversed by treating adults with a JH analog (30). Now we see that the insulin from the MNC regulates JH titer via insulin receptors upon the CA, and JH is the ultimate endocrine factor by which reduced insulin extend lifespan in response to gut NPF. Gut NPF modulates *Drosophila* aging through the integration of nutrient sensing, insulin signaling and juvenile hormone titer.

## Results

### Closed-type EEC secrete NPF in response to feeding

Previous studies used the intensity of peptide labeled with antibodies within gut EEC cells to infer when these cells secrete NPF (26, 27, 33). High stain intensity implies retention of the hormones, but high intensity can also arise if neuropeptides are actively secreted yet produced at an even greater rate. We therefore developed a method to directly quantify secreted NPF in adult hemolymph. We adapted an approach where a biotin ligase BirA*G3-ER (UAS-BirA*G3) is expressed in targeted cells (34). Adults with specific EEC drivers were fed biotin to label peptides produced in the endoplasmic reticulum of the target cells. Labeled neuropeptides from these cells could then be secreted into the hemolymph. We collected this hemolymph, captured the neuropeptides on streptavidin-coated plates, and quantified NPF with ELISA (Fig. 1A).

**Figure 1.**
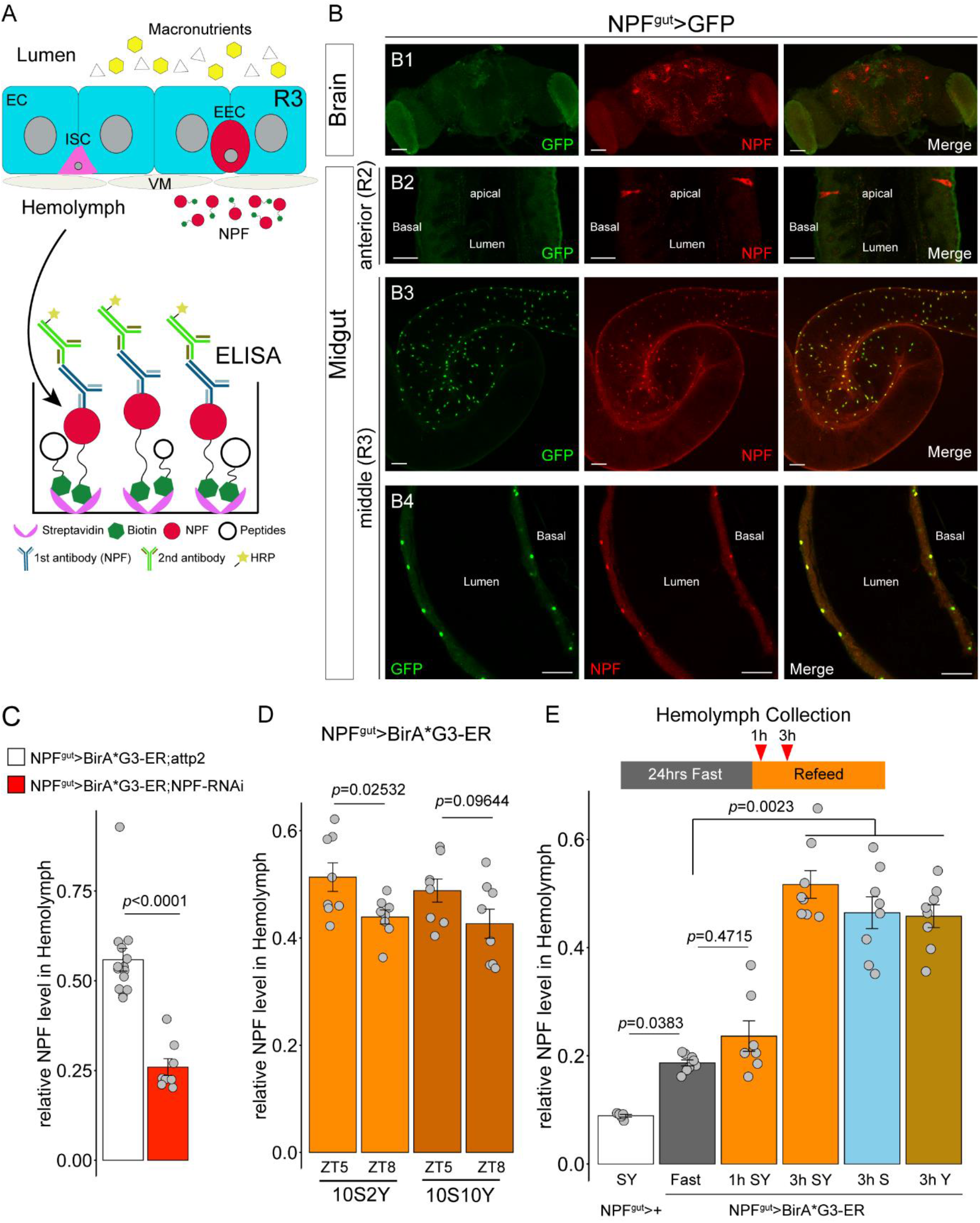
Closed-type EEC secrete NPF in response to feeding. (**A**) Design for ELISA to measure NPF secreted into hemolymph. Enterocyte (EC), Enteroendocrine cell (EEC), Intestinal stem cell (ISC), Visceral muscle (VM). (**B**) Gal4 specific to R3 closed-type EEC (NPF^gut^-Gal4: NPF-Gal4; nSyb-Gal80) drives UAS-GFP (green), co-labeled with ab-NPF (red). GFP is absent from NPF-positive neurons in the brain (**B1**) and NPF-positive open-type EEC of the anterior midgut R2 (**B2**). GFP is expressed in most NPF-positive closed-type EEC of the middle midgut R3 (**B3** and **B4**). Scale bars: **B1**-50 µm; **B2**-20 µm; **B3**- and **B4**- 50 µm. (**C**) NPF in hemolymph measured by ELISA when gut NPF was knocked down in closed-type EEC (NPF^gut^>BirA*G3-ER; NPF- RNAi) relative to NPF^gut^>BirA*G3-ER; attp2 (*t*-test). (**D**) NPF in hemolymph secreted from closed-type EEC from females maintained on low or high yeast diet at ZT 5 and ZT 8. (Two-way ANOVA with interaction: Diet-by-ZT, F=0.075, *p=*0.7861.) (**E**) NPF in hemolymph secreted from closed-type EEC from fasted females refed for 1- or 3-hours upon different diets. (One-way Kruskal-Wallis ANOVA.)

NPF^gut^-Gal4 (nSyb-Gal80; NPF-Gal4, see (26, 35)) drives UAS-transgenes in closed-type EEC of the midgut R3 region (Fig. 1B3, 4) but not in NPF-producing neurons of the brain or in EEC of the gut R2 (Fig. 1B1, 2; Fig. S1A). From the hemolymph of adult flies, we detect gut NPF tagged with biotin by driving UAS-BirA*G3 with NPF^gut^-Gal4, and this titer is reduced by simultaneously driving NPF-RNAi in the closed-type EEC (Fig. 1C; Fig. S1B). Dietary yeast concentration (2% or 10%) did not affect the hemolymph NPF titer, although hemolymph NPF was reduced at *zeitgeber* 8 (ZT 8) relative to earlier in the day (ZT 5) (Fig. 1D). To assess how feeding itself impacts NPF from closed-type EEC, flies were fasted for 24 hours and then fed a full (sugar + yeast), sugar-only or yeast-only diet. NPF in the hemolymph was equally elevated after refeeding across all diet types (Fig. 1E). These secretory responses were also observed with a calcium-dependent nuclear import reporter (CaLexA) (36, 37), in which GFP intensity, indicative of active secretion, was strong in EEC when flies were fed sugar or yeast (Fig. S1C). Thus, closed-type EEC are not sensitive to the concentration of dietary yeast but secrete NPF when refed sugar or yeast, and this level declines some hours after feeding. NPF from the closed type EECs may provide a satiety signal as suggested by Malita *et al*. (26) and by Gao *et al*. (33).

### Open-type EEC secrete NPF in response to dietary yeast concentration

NPF is also seen in open-type EEC of the midgut R2 region. Open-type EEC (14) have an apical extension to the gut lumen (Fig. 2A1). The Tkg-Gal4 construct drives expression in NPF-positive open-type EEC of R2 (Fig. 2A1, A2), and in a few closed-type EEC in R3 (Fig. 2A3), but not in NPF-producing neurons of the brain (27). Unlike the dietary response observed in closed-type EEC, 10% dietary yeast increased NPF secretion to hemolymph from open-type EEC (Fig. 2B). Elevated secretion was apparent at ZT5 but not at ZT8. Likewise, CaLexA signal in open-type EECs was elevated in flies fed 10% yeast (Fig. 2C, D). Notably, these secretory events occurred even while we saw high NPF staining intensity within these EECs (Fig 2C, E). Overall, high dietary protein promotes NPF secretion from open-type EEC (Fig. 2F).

**Figure 2.**
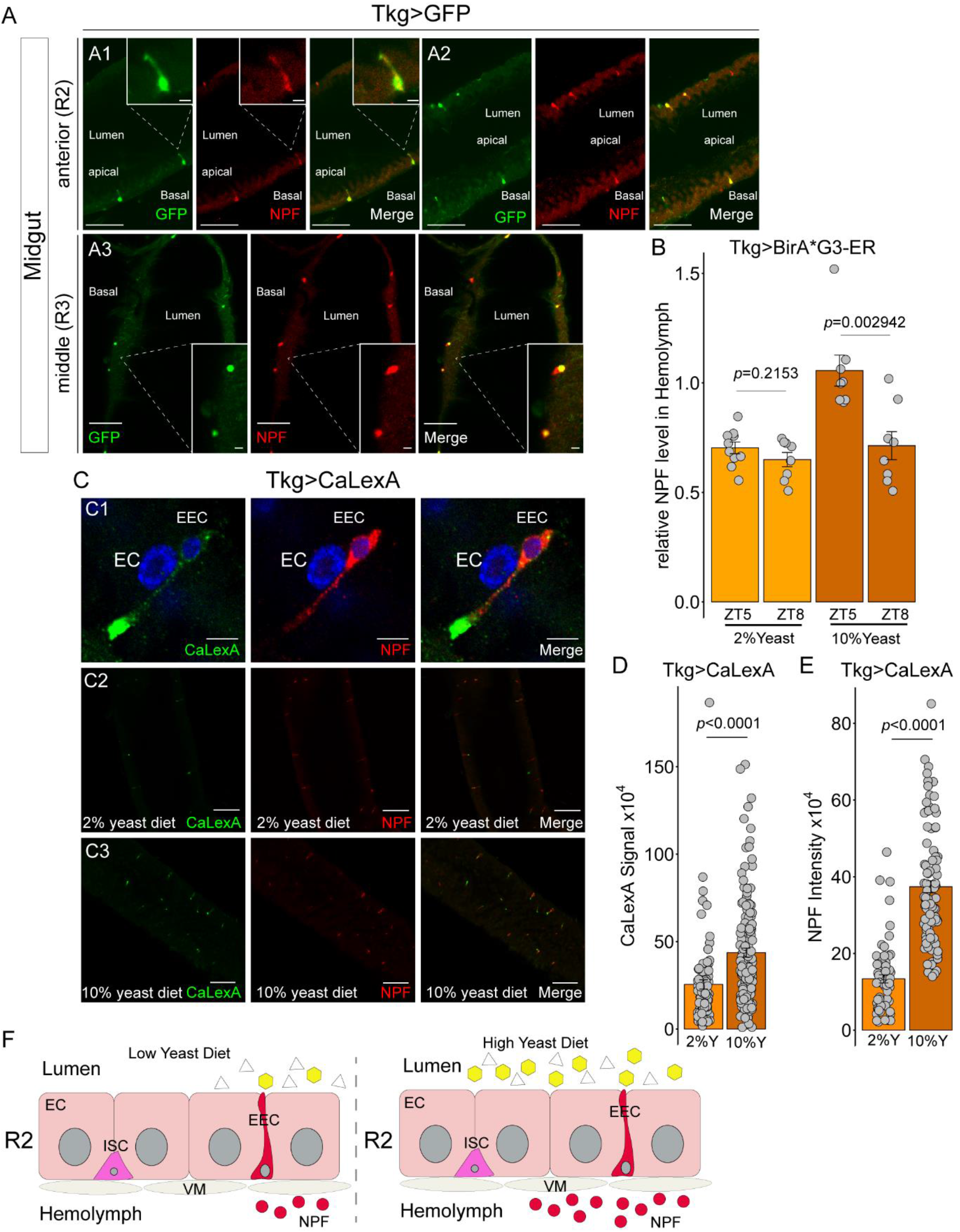
Open-type EEC secrete NPF in response to dietary yeast concentration. **(A)** Tkg-Gal4 drives in open-type EEC that produce NPF, visualized from UAS-GFP (green) and anti-NPF (red) in anterior R2 midgut (**A1**, **A2**) and some closed-type EEC in the middle R3 midgut (**A3**). (**B**) Biotin tagged NPF (Tkg- Gal4>BirA*G3-ER) in hemolymph from females maintained on low or high yeast diet, measured at ZT 5 and ZT 8. (Two-way ANOVA: Diet-by-ZT, F=8.329, *p=*0.0072.) (**C**) CaLexA activity (green) visualized at NPF-positive (red; nuclei: DAPI, blue) open-type EECs on low (**C2**) or high yeast (**C3**); quantified in (**D**, **E**) (Wilcoxon rank-sum tests). Scale bars: **A**-50 µm (5 µm); **C1**-5 µm; **C2-** and **C3**-50 µm. Enterocyte (EC), Enteroendocrine cell (EEC). (**F**) Summary for how dietary yeast impacts hemolymph NPF secreted from open-type EEC. Enterocyte (EC), Enteroendocrine cell (EEC), Intestinal stem cell (ISC), Visceral muscle (VM).

### Lifespan is modulated by NPF receptors at insulin producing cells of the brain

Yeast dietary restriction (DR) extends *Drosophila* longevity, and we now find that open-type EEC secrete NPF in response to the high dietary yeast. We therefore determined if limiting NPF from open-type EEC is required for diet restriction to extend lifespan. Knockdown of gut NPF with Tkg-Gal4 increased lifespan of adults upon high dietary yeast but not when adults were fed 2% yeast (Fig. 3A, 3B, S2A). Relative to wild type, depletion of gut NPF blunted the ability of DR to slow aging (genotype-by-diet interaction: *p*<0.0001). In contrast, closed-type EEC secrete NPF in response to feeding but do not respond to the concentration of dietary yeast. Accordingly, knockdown of NPF with NPF^gut^-Gal4 improved lifespan in flies maintained on both high and low dietary yeast (Fig. 3C, 3D, S2B). NPF from closed-type EEC modulates aging independent of dietary restriction (genotype-by-diet: *p=*0.1519).

**Figure 3.**
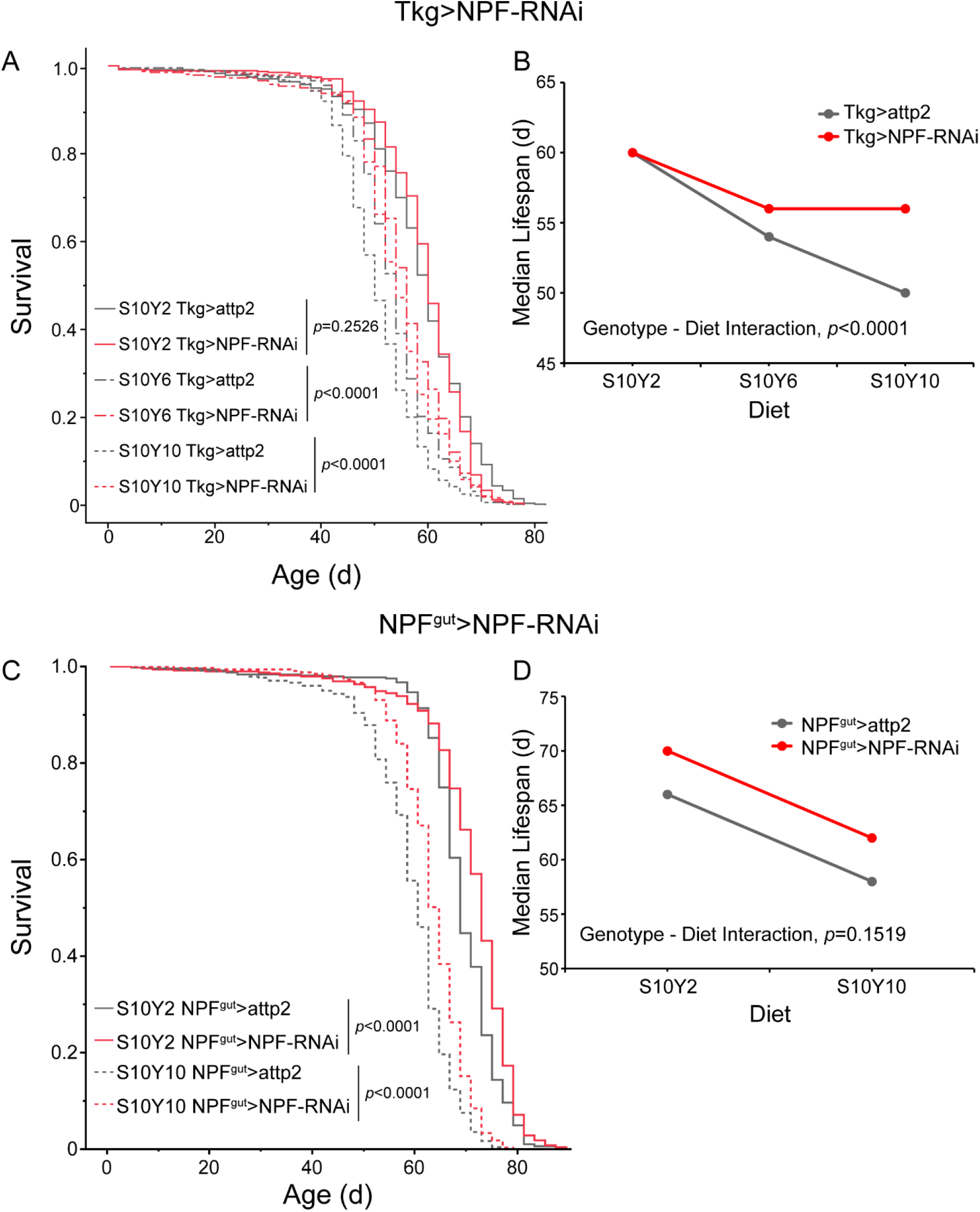
Depletion of gut NPF slows aging. (**A**) Survival of adult females maintained on diets with 10% sugar (S10) and yeast at either 2% (Y2) or 10% (Y10), while expressing NPF-RNAi in open-type EEC (Tkg-Gal4) relative to wild type. Pairwise survival analyses by log-rank tests. (**B**) Reaction plots of median lifespan of panel **A** cohorts. We observe significant genotype-by-diet interaction inferred from proportional hazard analysis: Genotype, *p*<0.0001; Diet, *p*<0.0001; Genotype-by-Diet, *p*<0.0001. (**C**) Survival was improved on both low and high yeast diet in females where NPF was depleted from closed-type EEC (NPF^gut^-Gal4). (**D**) Reaction plots of median lifespan of panel **C** cohorts. Knockdown of NPF equally increases lifespan on 2% and 10% yeast diet (Proportional hazard analysis: Genotype, *p*<0.0001; Diet, *p*<0.0001; Genotype-by-diet, *p=*0.1519). Further survival analysis statistics in **Table S1**.

### Lifespan is modulated by NPF receptors at insulin producing cells of the brain

Gut NPF secreted into hemolymph will act on tissues expressing the neuropeptide F receptor (NPFR) (26, 27, 35). To determine which tissue modulates lifespan in response to gut NPF, we measured survival when NPF receptors were depleted in ovaries (38), fat body, *corpora cardiaca* (CC, site of AKH synthesis) and brain median neurosecretory cells (MNC, site of insulin synthesis). Median lifespan was robustly extended (10 days), when NPF receptors were depleted in MNC (Fig. 4A, S3A), while knockdown of NPF in other tissues increased or decreased median lifespan no more than two days (Fig. 4B-D, supplemental data Table S1). Previous research found insulin secreted from MNC was reduced by knockdown of gut NPF or of MNC NPF receptors (27). We confirmed this observation (Fig. S4) and further found NPF secreted from the gut can locate at brain MNC neurons (Fig. S5). Gut-secreted NPF may thus act directly upon MNC NPF receptors to modulate insulin secretion. Given that altered insulin signaling slows *Drosophila* aging (30–32), reduced gut NPF is expected to extend lifespan because this impairs insulin secretion from the MNC.

**Figure 4.**
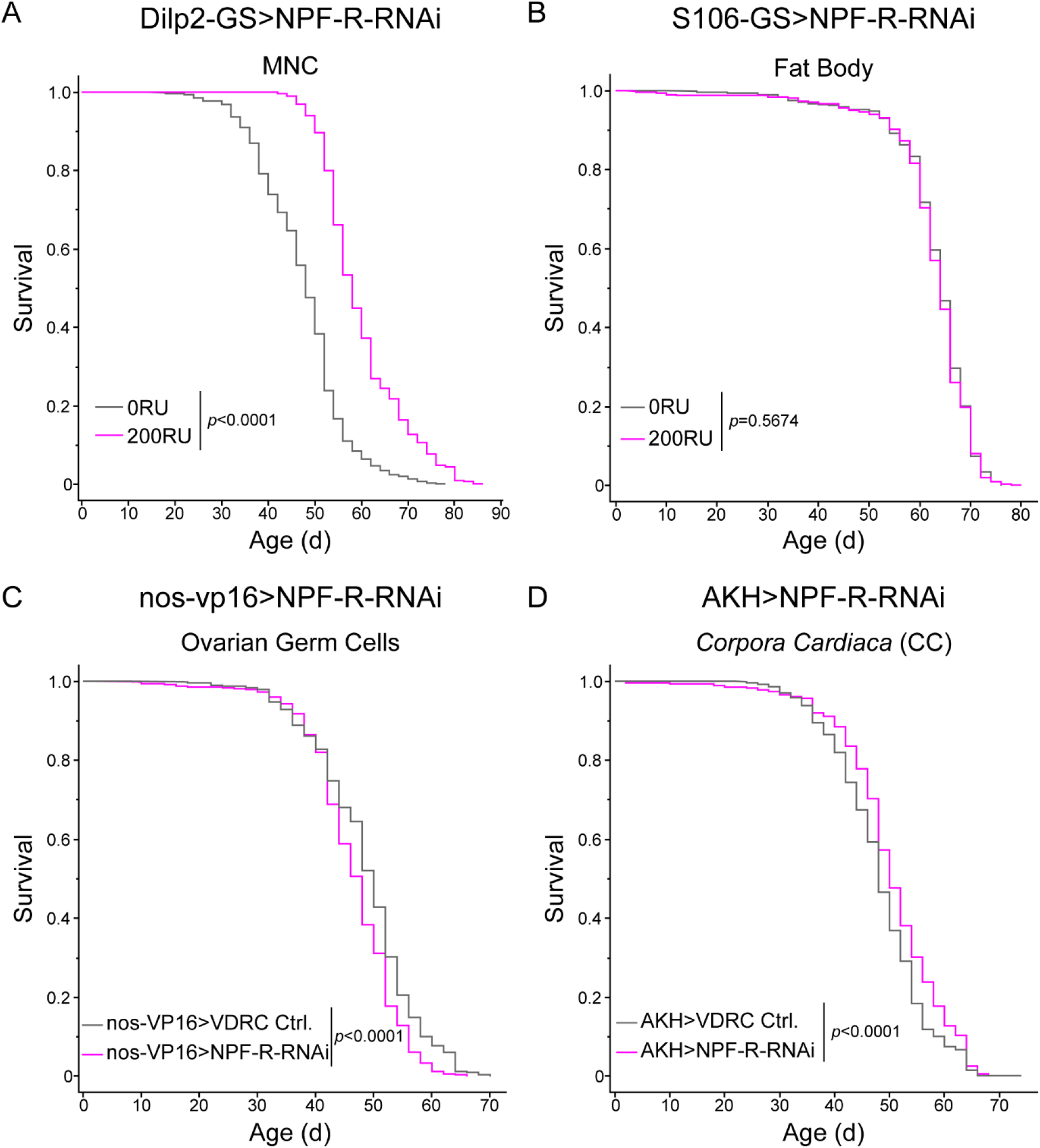
Knockdown of NPF receptors in the MNC extends lifespan. Female survival was robustly increased by *dilp2*-GS>UAS-*NPFR*-RNAi activated by RU486 relative to the same genotype without RU (log rank test *p* < 0.001). (**B-D**) Survival was minimally affected when NPF was depleted (UAS-NPF-RNAi) in: (**B**) Fat body; (**C**) Germline stem cells; (**D**) *Corpora cardiaca*. Further survival statistics in **Table S1**.

### The NPF-insulin axis regulates aging through control of juvenile hormone titer

Mutation of the *Drosophila* insulin receptor (dInr) prolongs longevity and suppresses JH production in the adult *corpora allata* (CA), an endocrine gland innervated by MNC (30). This extended longevity of *dInr* mutants was restored to wild type by treating adults with the JH analog methoprene (JHA), and subsequent work showed that ablation of the adult CA itself robustly extends lifespan (39). JH appears to be a master endocrine regulator of *Drosophila* aging. Accordingly, we determined if NPF regulates lifespan because the gut hormone controls JH titer.

We measured JH activity through its induction of Krüppel homolog 1 (*Kr-h1*), a target of the JH transcriptional complex Met/Gce (40). *Kr-h1* mRNA was reduced in flies when gut NPF was depleted by RNAi (Fig. 5A) and when NPF receptors at the MNC were depleted (Fig. 5B). Furthermore, we quantified the level of the circulating epoxidated JH in females via LC-MS/MS. Juvenile hormone III (JH III) and juvenile hormone III bisepoxide (JHB3) were reduced by depletion of NPF receptors at the MNC (Fig. 5C). To confirm that the loss of JH slows aging when we deplete gut NPF or MCN NPF receptors, these adults were treated with the JH analog methoprene. Methoprene reduced the survival of both genotypes to that of JHA treated wild type controls (Fig. 5D, E; Fig. S6). Gut NPF ultimately modulates aging through the action of juvenile hormone.

**Figure 5.**
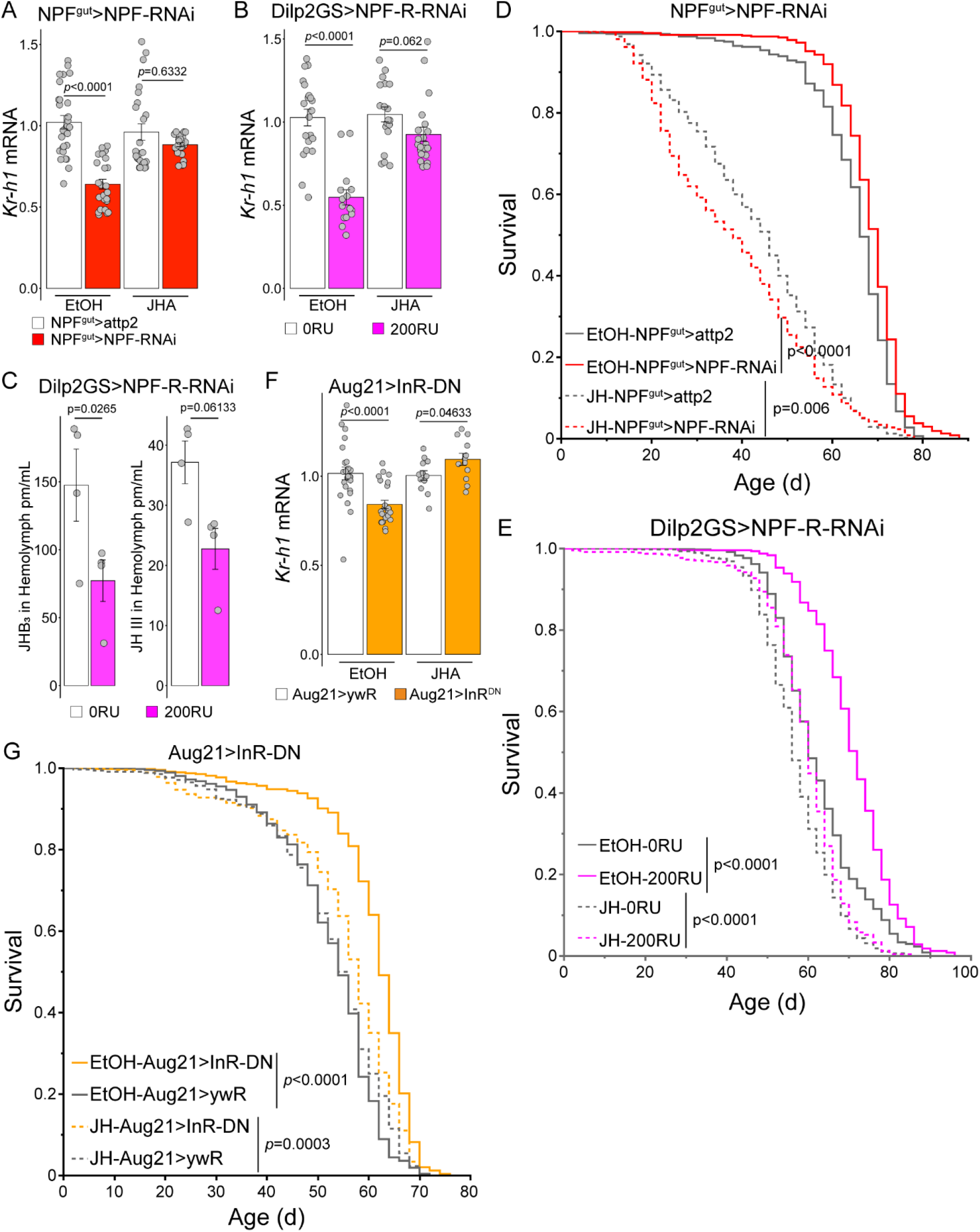
Juvenile hormone modulates longevity conferred by gut NPF. (**A**) Knockdown of gut NPF (NPF^gut^>NPF-RNAi) reduced expression of *Kr-h1*, where treatment with JHA restored *Kr-h1* mRNA (Two-way ANOVA: Genotype, F=39.204, *p*<0.0001; Treatment, F=6.482, *p=*0.0125; Genotype*Treatment, F=17.917, *p*<0.0001.) (**B**) Knockdown of NPF receptors in MNC (Dilp2GS>NPFR-RNAi) activated by RU reduced *Kr-h1* mRNA, where treatment with JHA restored *Kr-h1* mRNA to the level of wild type (Two-way ANOVA: RU, F=32.93, *p*<0.0001; Treatment, F=15.82, *p*<0.0001; RU*Treatment, F=14.62, *p=*0.0003.) (**C**) JHB3 and JH III in hemolymph is reduced by RU-induced knockdown of NPF receptors in MNC (Dilp2GS>NPFR-RNAi). (**D**) Knockdown of NPF in the gut extends lifespan (NPF^gut^>NPF-RNAi), where JHA reduces lifespan and eliminates the advantage conferred by NPF knockdown (Two-way Cox-proportional hazard analysis, treatment- by-genotype interaction, *p*=0.0002). (**E**) Knockdown of NPF receptor in the MNC extends lifespan (Dilp2GS>NPFR-RNAi), where JHA rescues lifespan to the level of wild type (Two-way Cox-proportional hazard analysis, treatment-by-RU interaction, *p*< 0.0001). (**F**) Inhibition of *dInr* the dominant-negative *dInr^DN^*expressed in the *corpora allata* (Aug21> dInr^DN^) reduced systemic *Kr-h1* mRNA, while treatment with JHA restored *Kr-h1* mRNA to wild type (Two-way ANOVA: Genotype, F=7.278, *p=*0.0088; Treatment, F=12.766, *p=*0.0007; Genotype-by-Treatment, F=15.416, *p=*0.0002.) (**G**) Inhibition of insulin receptors in the *corpora allata* extended lifespan, while treatment with JHA rescued this survival toward that of control genotype with and without JHA (Two-way Cox-proportional hazard analysis, treatment-by-RU interaction, p< 0.0001). Further survival statistics in **Table S1**.

JH is synthesized in the adult *corpora allata*. NPF from the gut could potentially modulate the CA directly, however the CA is not reported to contain NPF receptors (41), and NPFR-RNAi driven in the CA does not reduce systemic *Kr-h1* mRNA or the titer of JH (Fig. S7A). In contrast, the CA express insulin receptors (*dInr*) (42). To determine if insulin peptides from the MNC directly regulate JH, we inhibited insulin receptors in the CA by expressing a receptor dominant negative construct (Aug-21>*dInr*^DN^). This reduced systemic *Kr-h1* mRNA (Fig. 5F) and robustly increased lifespan (Fig. 5G; Fig. S7B). JHA treatment reduced the survival benefit of the Aug-21> *dInr*^DN^ adults (Fig. 5G). Potentially, this rescue could arise if methoprene induces insulin synthesis, but the JH analog did not elevate *Drosophila insulin-like peptide 2* (*dilp2*) mRNA (Fig. S7C). JH itself appears to regulate longevity downstream of insulin from the MNC.

## Discussion

### Multiple pathways modulate JH synthesis

Juvenile hormones are central regulators of insect development and life history where the biosynthesis of JH is concurrently influenced by factors such as developmental stage, nutrition, mating, circadian rhythms and temperature (43). To regulate the rate of JH synthesis in response to this complexity, the CA expresses receptors for multiple allatoregulatory factors (44). Ecdysis triggering hormone synthesized by the Inka cells located alongside the tracheal system stimulate JH biosynthesis. Likewise, although not in *Drosophila*, JH synthesis in the CA in many insects is stimulated by Allatotropin derived from neurosecretory cells of the brain. Allatostatic molecules that inhibit JH synthesis include Allatostatin C and Diuretic Hormone 31 produced by dorsal clock neurons and CA-projecting neurons. While the pathways involved in response to stage, mating, circadian rhythms and temperature have been recently described (45–48, 69), little is known about how nutrients are sensed in the gut to affect CA activity and in turn how this affects lifespan.

### Gut NPF controls longevity in response to diet

Enteroendocrine cells (EEC) are scattered throughout the epithelium of the *Drosophila* gastrointestinal tract. Here we find that limiting dietary yeast, which is associated with extended lifespan (49, 50), reduces the secretion of NPF from open EEC into the hemolymph. Previous work with *Drosophila* showed that survival was decreased when the non-specific secretion from all NPF-positive cells was increased (51). Here we show that specific knockdown of gut NPF is sufficient to *extend* adult *Drosophila* survival. Notably, NPF from open-type EEC of the midgut R2 region modulates how dietary (yeast) restriction slows aging. NPF is also secreted from closed-type EEC, but blunting NPF from these cells extends lifespan equally on low and high protein diets. Overall, gut NPF can regulate fly aging in response to dietary protein but also independently of diet quality.

### An NPF-insulin-JH axis controls aging

Gut NPF secreted into the hemolymph may act on several tissues. Yoshinari (27) previously showed that knockdown of gut NFP reduces insulin secreted from MNC of the brain. Here we biotinylated neuropeptides produced in gut EEC and tracked secreted NPF to the MNC. Lifespan was increased when NPF receptors at the MNC were depleted. This receptor knockdown reduced insulin peptide mRNA, as seen in Yoshinari (27). Many studies suggest altered insulin/IGF signaling non-autonomously slows aging but the underlying mechanisms are poorly understood. We previously suggested insulin signaling affects *Drosophila* aging through control of juvenile hormone titer (30, 52), and now show that gut NPF and NPF receptors at the MNC regulate aging because they control JH titer in response to insulin-like peptide.

Our data demonstrate that aging is modulated by interorgan communication between the gut, the brain, and the *corpora allata*, and insulin signaling affects aging because *Drosophila* insulin is an allatotropric hormone, increasing JH biosynthesis. This outcome suggests a model that coalesces previous observations (Fig. 6). The adult *corpora allata* produces juvenile hormone, which modulates a range of adult traits including egg production (46, 53), behavior (48), intestinal remodeling (54) and lifespan (39). The synthesis of JH by the CA is regulated through multiple allatostatic and allatotropic factors (41, 42, 44). Although we have long known that altered insulin likewise mediates JH production in *Drosophila* (30, 52), here we show this control is achieved directly at the CA through insulin receptors that respond to insulin produced by MNC of the brain. Insulin from the MNC is regulated by many inputs (serotonin, octopamine, GABA, short neuropeptide F (sNPF), corazonin and tachykinin-related peptide) (55), including by gut NPF as first reported by Yoshinari (27). Each therefore has the potential to impact adult JH production. Here we find that NPF produced by the gut can regulate JH produced in the CA, where this connection is relayed by brain medial neurosecretory cells.

**Figure 6.**
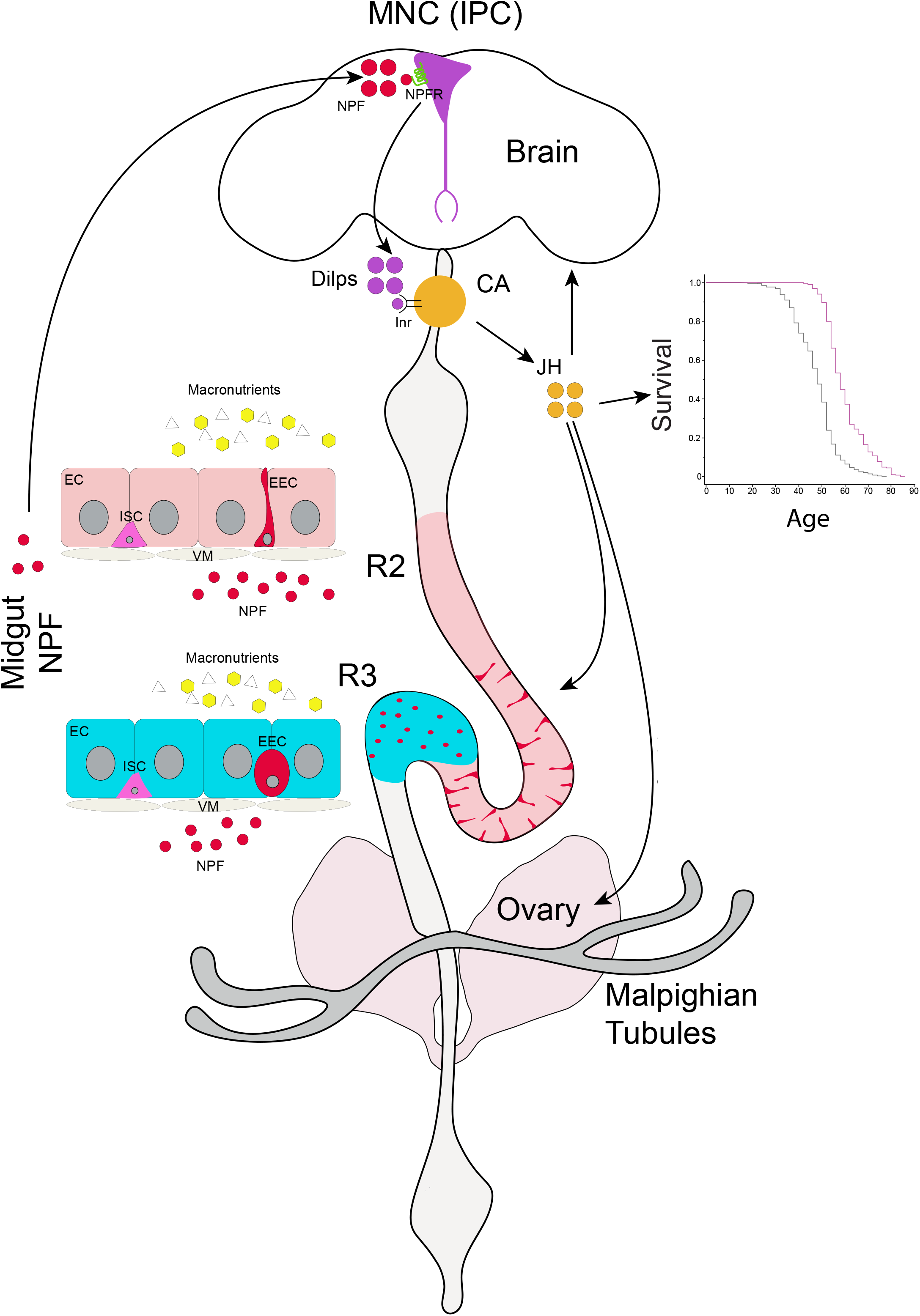
Model of interorgan communication between gut-brain-*corpora allata* in *Drosophila melanogaster*. The *Drosophila* midgut contains enterocyte cells (EC), intestinal stem cells (ISC) and enteroendocrine cells (EEC). Dietary macronutrients (yeast and sugar) induce EEC to secrete Neuropeptide F (NPF) that crosses the ventral muscles (VM) into the hemolymph. Open-type EEC of gut region R2 secrete NPF in response to the concentration of dietary yeast. Closed-type EEC of gut region R3 secrete NPF in response to feeding. Gut NPF enters the brain where it can localize at median neurosecretory cells (MNC) that produce insulin (insulin producing cells, IPC). NPF receptors at the MNC will be activated by the gut-produced NPF and stimulate these neurons to secrete *Drosophila* insulin-like peptides (Dilps). Dilps stimulate insulin-like receptors (Inr) at the *corpora allata* (CA), which induces the synthesis and release of juvenile hormone (JH). JH is highly pleiotropic, acting at many tissues to affect a range of adult phenotypes including behavior (brain), gut remodeling, egg production (ovary) and adult survival.

### A potentially conserved gut-brain-endocrine axis controls aging

How does JH ultimately modulate the aging process? JH is transiently elevated in newly eclosed adult *Drosophila* (53) where it controls the final development of the adult gut, fat body, and gonads (54). Adult JH likewise promotes intestinal stem cells (ISC) division during gut regeneration (56). JH is repressed during adult *Drosophila* reproductive diapause, a state demonstrated to slow aging (57). To further understand how JH affects *Drosophila* aging, Yamamoto *et al*. (39) described transcriptional profiles from adults where ablation of the *corpora allata* extended lifespan. Slow aging was associated with an increase in the amino acid storage protein larval serum protein 1 (LSP1), fatty acid binding protein 1 (FBP1) and the odorant binding protein Obp99b, as well as downregulation of proteases such as Jon25Bii. Most recently, Kim *et al*. (58) reported that a transient increase in JH titer in early adulthood disrupts protein homeostasis (proteostasis) and decreases lifespan.

Mammals have no direct analog of JH, but the insect CA and mammalian hypothalamic-pituitary system may represent parallel endocrine architectures, which suggest growth hormone or thyroid hormone may modulate aging as we see with JH (59). The *Drosophila* incretin NPF resembles mammalian NPY, peptide YY (PYY) and pancreatic polypeptide (PP). Although NPY is a hypothalamic hormone, it is also produced in the mammalian gut where it influences gastrointestinal motility, electrolyte secretion and immune function (60). Likewise, PYY is produced in small intestine enteroendocrine L-cells where it is co-released with the incretin glucagon-like peptide 1 (GLP-1) (61). Both NPY and PYY are proposed to influence mammalian age associated diseases and longevity (62–64), while PYY is thought to act directly on Y1 receptors in the pancreas as well as in the central nervous system to inhibit insulin secretion (65, 66). Here we find reduced gut NPF is sufficient to slow *Drosophila* aging. This result prompts us to ask whether secretion of gut neuropeptides from EEC might impact human aging, and whether treatment with analogs of gut incretin hormones may alter the aging trajectory independent of their immediate metabolic benefits.

## Material and Methods

### Fly husbandry and stocks

Stocks were reared on standard *Drosophila* medium containing 11% sugar, 2.5% yeast, 5.2% 0.8% agar (w/v in 100 mL water) with 0.2% Tegosept (methyl 4- hydroxybenzoate, Sigma), and maintained at 25°C, 40% RH and 12:12 light-dark cycle (LD). RU486 (mifepristone, Sigma) used to activate GeneSwitch-Gal4 was dissolved in ethanol to a concentration of 200 μM and added to cooked food.

Fly lines were backcrossed to *yw* background for at least 8 generations. Stocks: NPF- Gal4 (BDSC#25682), Tkg-Gal4 (provided by N. Perrimon, HHMI), VP16-nos-Gal4 (BDSC#4937), Dilp2-GeneSwitch-Gal4 (70), S106-GeneSwitch-Gal4 (67), AKH-Gal4 (BDSC#25683), Aug21-Gal4 (BDSC#30137), nsyb-Gal80 (35) (provided by S. Pletcher, University of Michigan), UAS-NPF-RNAi (BDSC#27237), UAS-attp2 (BDSC#36303), UAS-InR^DN^ (BDSC#8253), UAS-NPF-R-RNAi (VDRC KK107663), UAS-VDRC-control (VDRC 60100), UAS-20xUAS-IVS-mCD8::GFP (BDSC#32194), UAS-CaLexA (BDSC#66542), UAS-BirA*G3-ER-Myc (34) (provided by N. Perrimon, HHMI).

### Demography and survival analysis

Newly eclosed adults were transferred to fresh standard food in bottles to mate for 48 hours at 25°C, 40% RH and 12:12 LD cycle. 48 hours old female adults were sorted with light CO2 and then pooled in 1L demography cages maintained at 25°C, 40%RH and 12:12 LD cycle with an initial density of 125 adults per cage. Four cages were initiated per genotype. Food vials of demography cages contained 10% sugar, 5.2% cornmeal, 0.8% agar and 2% or 10 % yeast for 2%- or 10%-yeast diet, respectively. Food vials were changed daily for the first 45 days and every two days thereafter. Dead flies were removed from each cage and counted every two days. Treatment with juvenile hormone analog (Methoprene; Sigma PESTANAL, racemic mixture) followed the methods of Yamamoto (39): 10µl of 32.2 µM methoprene in ethanol was applied to a cotton bud fixed within each demography cage and replenished twice weekly.

### Immunostaining and imaging

Tissues of adult flies were dissected in phosphate-buffered saline (PBS) and fixed in 4% paraformaldehyde (PFA) in PBS at room temperature with gentle shaking for 45 min (gut) or 90 min (adult brain). After six washes with PBS+0.3% Triton-X, samples were blocked in PBS+0.3% Triton-X with 5% normal goat serum (NGS, Thermo Fisher) at room temperature for 2 hours. After six further washes with PBS+0.3% Triton-X, samples were incubated in primary antibody dilutions in PBS+0.3% Triton-X with 5% NGS at 4°C overnight. Primary antibodies included: anti-NPF 1:1000 (RayBiotech RB-19-0001), anti-GFP 1:2000 (Abcam ab13970), anti-P-Glycoprotein (C219) 1:100 (Invitrogen MA1-26528). Washed samples (6X) were incubated in secondary antibody dilutions and additional staining reagents in PBS+0.3% Triton-X with 5% NGS at room temperature for 2 hours. Secondary antibodies and reagents included: Goat anti-Chicken IgY 488, 1:200 (Invitrogen A11039), Goat anti-Mouse IgG 488 (Invitrogen A11029), Goat anti-Rabbit IgG 555 (Invitrogen A21428), anti-Streptavidin 647 (Invitrogen S32357), DAPI, 0.5µg/mL (Sigma). Samples were mounted in 80% glycerol. Images were acquired with a Zeiss LSM 800 Confocal Microscope and analyzed using ImageJ software. Total fluorescence intensity of identified EECs in confocal sections was measured using the 3D Object counter of ImageJ.

### Biotin-containing fly food and *Drosophila* culture

Biotin-containing food was prepared following Droujinine (34). 18 mM biotin stock was made by dissolving solid biotin (Sigma B4639) in water, adjusting the final pH to 7.2. Food with biotin at a final concentration of 100 μM was prepared, dried overnight and stored at 4 °C. EEC-Gal4 driver lines were crossed to UAS-BirA*G3-ER. Progeny were allowed to mate for 2 days at 25°C on standard food without biotin. Females were switched to 29°C for 2 days on standard food, and then transferred to food with 100 μM biotin while at 29 °C for 6 days. Fresh food was provided daily.

### Hemolymph Collection

Hemolymph was processed from biological replicates, each made from a pool of 20 flies. Flies were punctured at the thorax with a tungsten needle (Fine Science Tools) and transferred to a 0.5 mL tube where the bottom contained a 25-gauge hole. To collect hemolymph for ELISA, the 0.5 mL tube was inserted into 1.5 mL low-protein binding tube and the two-tube assembly centrifuged at 9000rpm for 5 min at 4°C. The supernatant was kept on ice and biotinylated NPF was pulled down on 96-well streptavidin-coated plates and quantified by ELISA (see below). To collect hemolymph for JH measurement, the 0.5 mL tube with flies was inserted into a siliconized glass vial (Fisher Scientific 03-395G), with the assembly centrifuged at 9000 rpm for 5 min at 4°C. The supernatant was kept on ice and hemolymph JH was extracted and measured by LC-MS/MS (see below).

### Protein lysate preparation

For lysis buffer, RIPA (Pierce) was supplemented with PMSF (Sigma) PhosSTOP (Roch) and cOmplete EDTA-free Tablet (Roch). For each biological replicate, 10 dissected brains were placed in lysis buffer with beads and homogenized with a TissueLyser (Retsch) in three cycles of 2 min. Samples were centrifuged for 15 min at 4°C. Collected supernatant was assayed for total protein concentrations (BCA protein assay kit, Pierce) and biotinylated gut-NPF was pulled down with streptavidin-coated plated and quantified by ELISA (see below).

### Streptavidin plate pulldown and Enzyme-linked immunosorbent assay (ELISA)

For each biological replicate of hemolymph or brain proteins, 1 μL of hemolymph or 2 μg of protein lysate was diluted into 200 μL of PBS + 0.05% Tween20. 100 μL of samples was pipetted into each well of a 96-well streptavidin-coated plate (Thermo Fischer 15125) and incubated on a shaker at 4°C overnight. Incubated plates were washed six times with PBS + 0.05% Tween20 and then exposed to anti-NPF antibody (1:1000) for 3 hours. After six washes, samples were incubated with HRP-conjugated secondary antibody at 1∶5000 (Jackson Immuno Research 111-035-144) for 2 hours. After six final washes, samples were treated with TMB solution (Cell signaling 7004) for 30 min until terminated with Stop solution (Cell signaling 7002). Absorbance was recorded at 450nm using a SpectraMax M5 (Molecular Devices LLC).

### JH extraction and measurement

5 μL hemolymph of each biological replicate was transferred into pre-chilled siliconized glass vials (Fisher Scientific 03-395G) on ice with a siliconized glass pipette and supplemented with 150 μL of chilled PBS from the same tip. 10 μL of 6.25 pg/μL JH III-D3 (Toronto Research Chemicals, #E589402) in acetonitrile was added into each sample along with 600 μL of hexane. Samples were vortexed for 1 min and centrifuged for 5 min at 4500 rpm. 500 μL of the organic phase was transferred to a fresh silanized glass vial. JH titers were measured using LC-MS/MS as described in Ramirez *et al*. (68).

### Quantitative RT-PCR

Total mRNA was extracted from 5 biological replicates of 8 females (once-mated, aged 9 d) for each genotype using Trizol reagent (Invitrogen 15596018) and Direct-zol RNA Miniprep Plus Kits (R2072). RNA yield was determined with a NanoDrop ND-1000 Spectrophotometer. cDNA was synthesized using iScript (Bio-Rad 1708890), and RT-qPCR performed by QuantStudio 3 (Applied Biosystems) with SYBR Green PCR Master Mix (Applied Biosystems 4309155). Relative mRNA abundance of each gene was normalized to *rp49*; see Table S2 for primer sequences.

## Statistical analysis

Data are presented as mean ± SEM from independent biological replicates. For data confirmed to be normally distributed, comparisons were based on Student’s *t*-tests or one-way ANOVA with post hoc multiple-testing, and Wilcoxon rank-sum or one-way Kruskal-Wallis ANOVA were used to evaluate non-normally distributed data. Two-way ANOVA was performed to determine the interaction between JHA treatment and manipulated genes, or diets and ZT. Survival analysis was conducted in JMP Pro 16 software with combined data from replicate cages. Log-Rank tests provided pairwise comparison of survival distributions. Proportional hazard survival analysis was used to infer if lifespan was affected by the interaction between JHA treatment and manipulated genes.

## Acknowledgments

Work reported in this paper was supported by funding to JC and MT from the National Institutes of Health (R01AG059563 and R37 AG024360), an NIH-NIAID R21 award to F.G.N. (R21AI167849) and a project 22-21244S from the Czech Science Foundation, Czech Republic, to MN. We thank our technical assistance at Brown University by Aleksandra Norton and Ilsa Dontes. We thank Lilian V. Tose from FIU for MS support.

## Supplemental Figures and Tables

**Figure S1.**
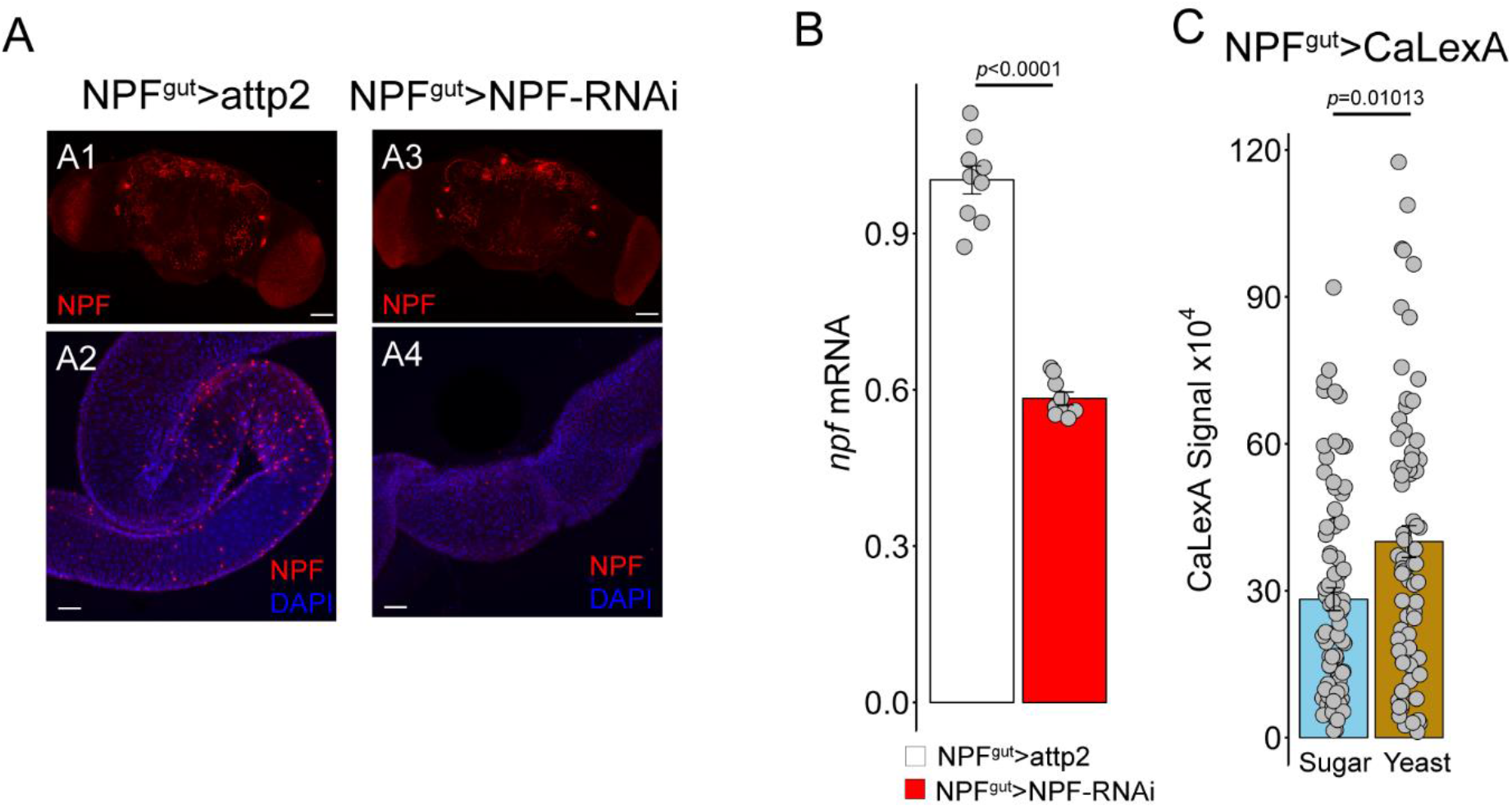
NPF^gut^-Gal4>NPF-RNAi knocks down NPF in EEC without affecting NPF producing cells of the brain. Closed type EEC show a global secretory signal when refed ar or yeast after fasting. **(A)** NPF (red) expressed in normal brain (**A1**) and gut (**A2**). Knockdown of NPF by RNAi depleted NPF in the gut (**A4**) without affecting NPF (red) in the brain (**A3**). In panels A: NPF labeled by antibody (red), nuclei by DAPI (blue). Scale bar, A:50 µm. **(B)** Relative *npf* mRNA in the total gut when NPF was knocked down in the R3 by PF^gut^>RNAi. *t-test* p<0.0001. NPF-RNAi was effective. **(C)** CaLexA signal (calcium dependent GFP) in closed-type EECs of R3 measures overall secretion from the cells. GFP from multiple cells from each of several independent midguts were quantified when adults were fasted and then refed either sugar-only or yeast-only diets.

**Figure S2.**
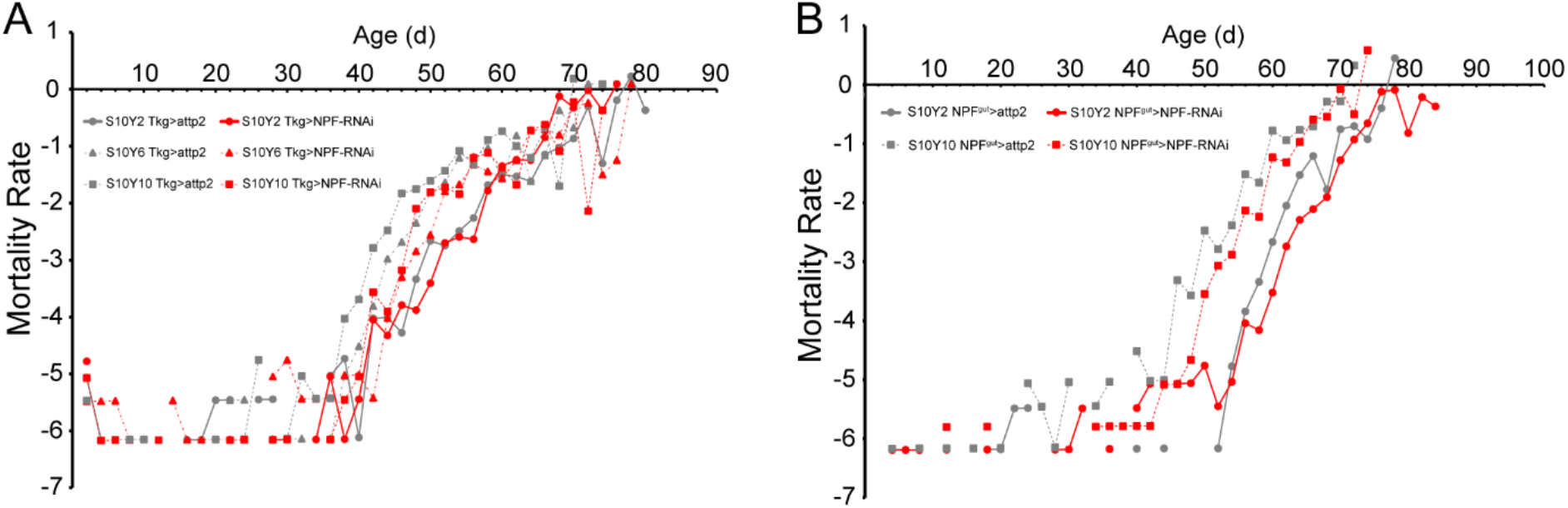
Mortality rate of cohorts in survival plots of Figure 3. The mortality plots show when demographic aging is affected by knockdown of NPF receptors in specific tissues. **(A)** NPF depleted by RNAi in open type EEC extends life on high yeast diets (S10Y6 and Y10) because this consistently reduces the progressive increase of mortality rate with age. At dietary yeast (S10Y2), mortality rate is similar across all ages among the control and NPF knockdown cohorts. **(B)** NPF depleted by RNAi in closed type EEC extends life on high yeast (S10Y10) and yeast diet (S10Y2) because knock down consistently reduces the progressive increase of mortality rate with age in both diet environments.

**Figure S3.**
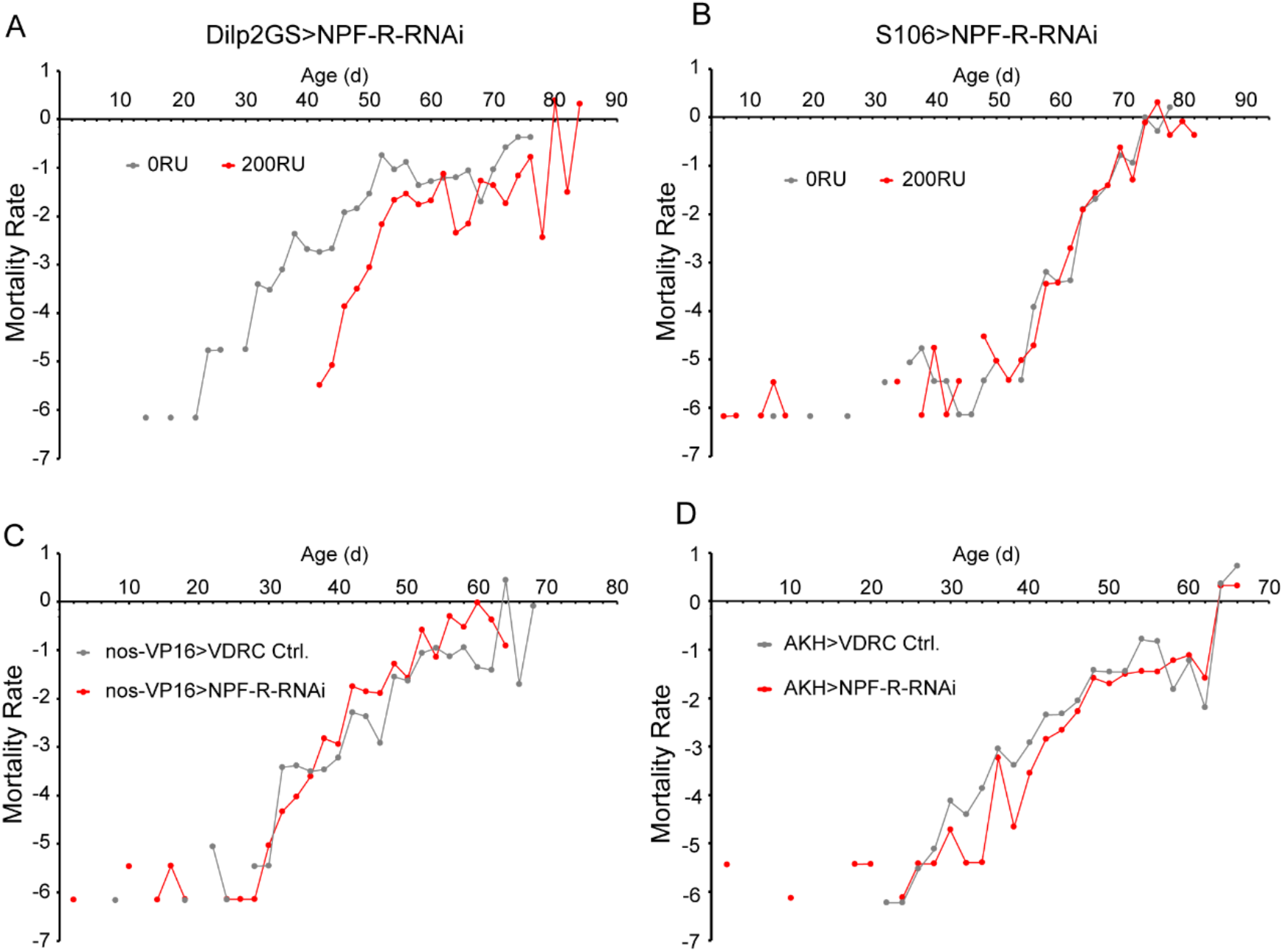
Mortality rate of cohorts in survival plots of Figure 4. Mortality plots demonstrate when demographic aging is affected knock down of NPF receptors in cific tissues. **(A)** Knock down of NPFR in MNC (A) consistently reduces the progressive increase of tality rate with age. RNAi is induced by RU486 acting upon dilp2(gene switch-gal4). **(B)** Knock down of NPFR in abdominal fat body does not affect mortality rate. RNAi is ced by RU486 acting upon S106(gene switch-gal4). **(C)** Knock down of NPFR in germline stem cells (nos-VP16) in backcrossed offspring tive to backcrossed VDRC control stock. Knockdown slightly increases mortality rate age 35 days onward, accounting for a modest reduction in median lifespan among cohorts seen in Figure 4C. **(D)** Knock down of NPFR in the *corpora cardiaca* modestly and consistently reduced tality rate. Offspring of both cohorts derived from back-crossed parentals, accounting he modest increase in median lifespan of the knockdown adults.

**Figure S4.**
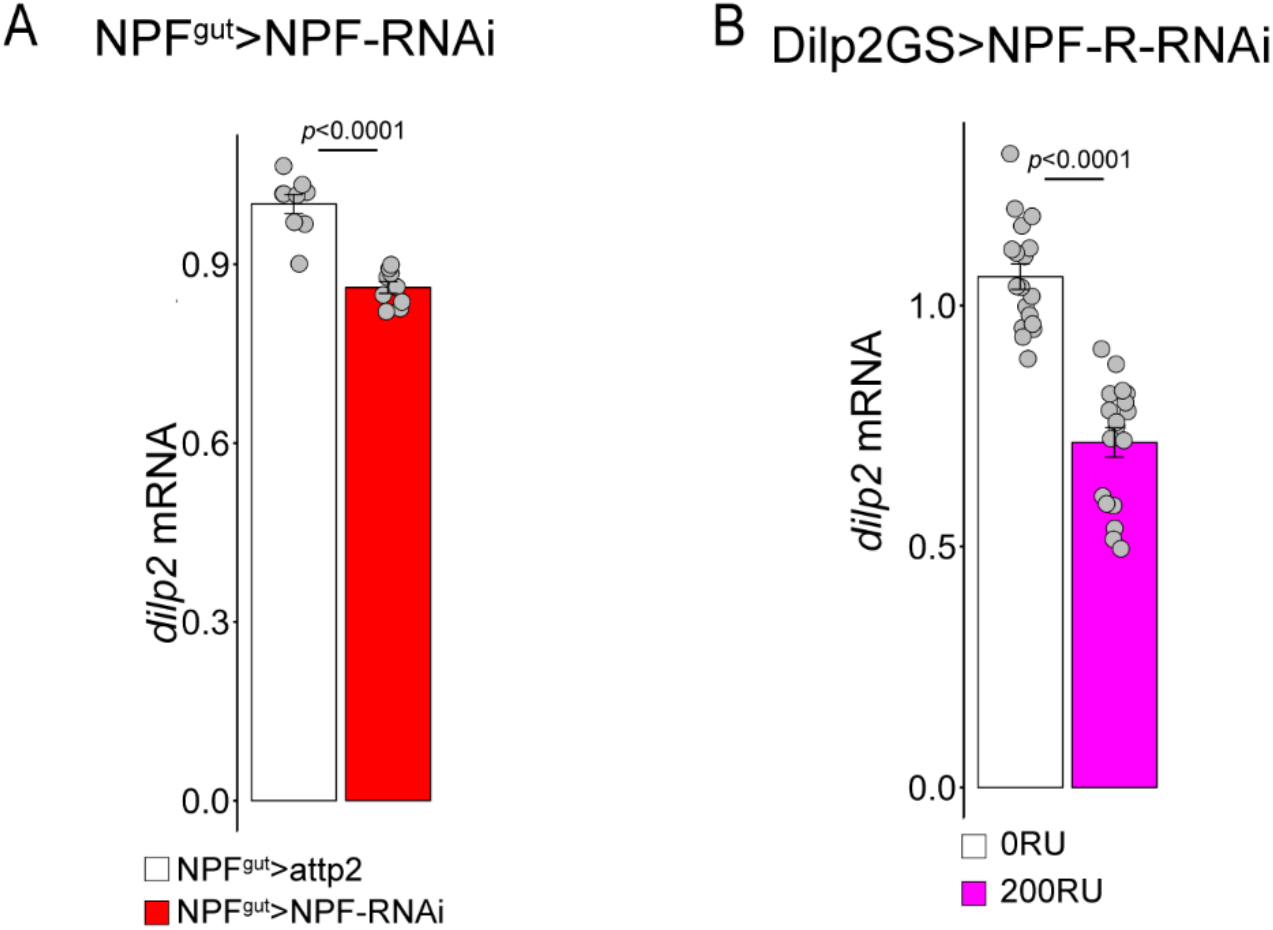
dilp2 mRNA is reduced when NPF is depleted from gut R3 EEC and when NPFR is depleted from insulin producing MNC of the brain. **(A)** *dilp2* mRNA measured from whole heads is reduced by knock-down of gut NPF by RNAi st p<0.0001). Offspring were tested from back-crossed parental genotypes. **(B)** *dilp2* mRNA measured from whole heads is reduced by knock-down of NPF receptors at MNC (t-test p<0.0001). RU486 activates gene switch.

**Figure S5.**
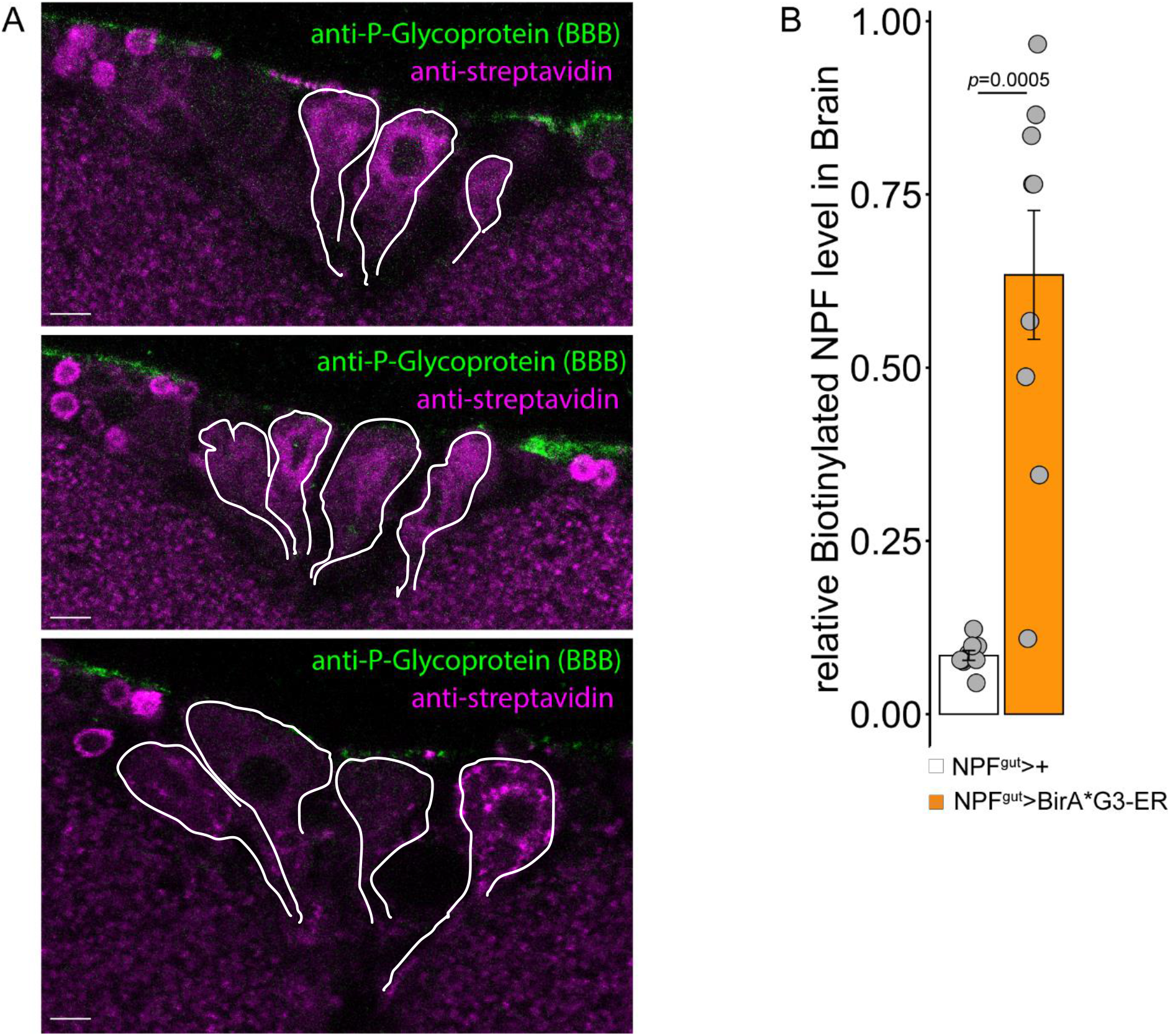
NPF biotin tagged and secreted from gut EEC is detected at the MNC within the brain. **(A)** Confocal images of MNC from NPFgut>BirA*G3-ER flies after feeding with biotin. Anti-streptavidin (purple) is enriched in cells of the MNC, identified here based on cell anatomy and location (outlined in white). The blood brain barrier is marked (green) by anti-P-Glycoprotein. Scale bar: 50 µm. **(B)** NPF from the gut tagged with biotin is strongly enriched in the whole brain from NPF^gut^>BirA*G3-ER flies after feeding with biotin, relative to genetic controls also fed biotin. NPF measured by ELISA from brain tissue after streptavidin pull-down. Wilcoxon rank sum test p=0.0005

**Figure S6.**
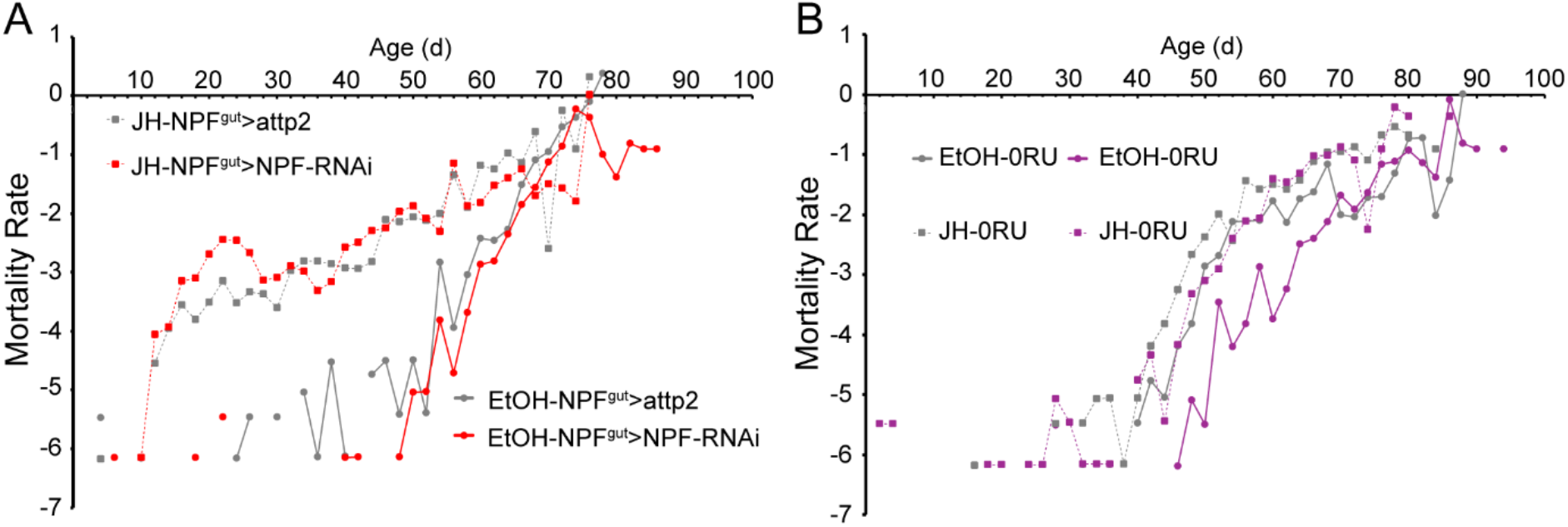
Mortality rate of cohorts corresponding to survival plots of Figure 5 D, E. **(A)** Knock down of gut NPF (closed EEC) modestly, consistently reduces mortality rate when treated with vehicle (ETOH) relative to control treated with ETOH (solid lines). JH treatment elevated mortality for both knockdown and control, and eliminated the mortality benefit of the knockdown relative to the control. **(B)** Knockdown of the NPFR in insulin producing MNC strongly reduced mortality rate in a cohort treated with vehicle relative to control treated with ETOH. JH treatment restored the knock down cohort mortality rate to the age-dependent trajectory of the control cohort, also treated with JH. The JH rescued the demographic aging of the NPFR knockdown.

**Figure S7.**
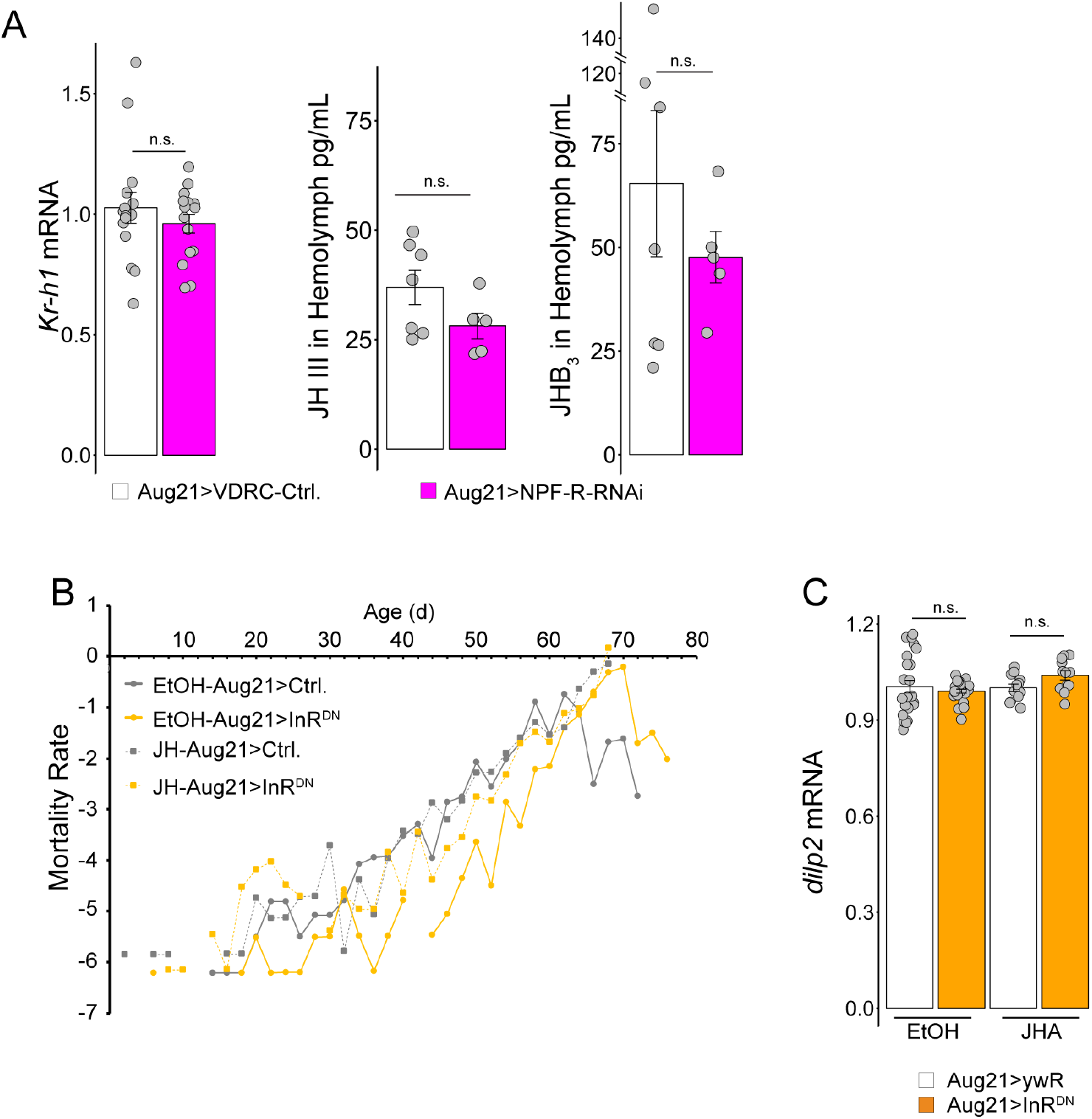
The dInr in the *corpora allata* modulates the mortality rate without affecting Dilp2 expressed in the brain, whereas the *corpora allata* do not express functional NPF receptors. **(A)** Targeting NPF receptor mRNA in the *corpora allata* did not affect JH titer measured indirectly by expression of JH responsive *Kr-h1* expression (*t-test* p=0.3871), or JH III and JHB3 as measured directly from hemolymph by LC-MS/MS. These data are consistent with absence of NPFR in the CA. **(B)** Mortality rate of cohorts corresponding to survival plot of Figure 5 F. Dominant negative inhibition of Inr in the CA consistently reduces mortality rate, and JH treatment rescues this reduced demographic aging to the mortality pattern of genetic controls. **(C)** Repressing dInR in the c*orpora allata* does not reduce *dilp2* mRNA expressed in the brain, nor does JH treatment. Inr at the CA does not regulate aging through feedback upon DILPs. Two-way ANOVA: Genotype, F=0.057, p=0.812; Treatment, F=2.043, p=0.158; Genotype*Treatment, F=2.718, p=0.104

**Table S1.** Survival analysis.

| Fig. | Genotype | Diet | Sample size | Mean | Mean s.e. | Median (d) | Lower 95% | Upper 95% | Test | $\chi^2$ | p |
| --- | --- | --- | --- | --- | --- | --- | --- | --- | --- | --- | --- |
| 3A | ywR/ywR;+;Tkg-ga4/UAS-attp2 | 10% sugar, 2% yeast | 477 | 58.3438 | 0.52147 | 60 | 58 | 60 |  |  |  |
| 3A | ywR/ywR;+;Tkg-ga4/UAS-NPF-RNAi | 10% sugar, 2% yeast | 477 | 59.1279 | 0.43513 | 60 | 60 | 62 | Log-Rank | 1.309 | 0.2526 |
| 3A | ywR/ywR;+;Tkg-ga4/UAS-attp2 | 10% sugar, 6% yeast | 476 | 53.2395 | 0.41353 | 54 | 52 | 54 |  |  |  |
| 3A | ywR/ywR;+;Tkg-ga4/UAS-NPF-RNAi | 10% sugar, 6% yeast | 482 | 54.9336 | 0.4965 | 56 | 54 | 56 | Log-Rank | 18.7698 | <0.0001 |
| 3A | ywR/ywR;+;Tkg-ga4/UAS-attp2 | 10% sugar, 10% yeast | 474 | 50.2954 | 0.41715 | 50 | 50 | 52 |  |  |  |
| 3A | ywR/ywR;+;Tkg-ga4/UAS-NPF-RNAi | 10% sugar, 10% yeast | 480 | 54.1958 | 0.43335 | 54 | 54 | 56 | Log-Rank | 50.8741 | <0.0001 |
| Term |  | Estimate | Std Error | Lower 95% | Upper 95% |  |  |  | Proportional hazard test |  |  |
| Diet[2% Yeast] | | -0.4122076 | 0.027001 | -0.465352 | -0.359496 | | | | Interaction | $\chi^2$ | p |
| Diet[6% Yeast] |  | 0.088491 | 0.026473 | 0.0363778 | 0.140162 |  |  |  | Genotype | 33.9346964 | <0.0001 |
| Genotype[Tkg>attp2] |  | 0.109426 | 0.018763 | 0.0726454 | 0.1462032 |  |  |  | Diet | 264.559461 | <0.0001 |
| Genotype[Tkg>attp2]*Diet[2% Yeast] |  | -0.1319642 | 0.026607 | -0.184124 | -0.079811 |  |  |  | Genotype*Diet | 29.8141588 | <0.0001 |
| Genotype[Tkg>attp2]*Diet[6% Yeast] |  | 0.012991 | 0.026437 | -0.038835 | 0.0648128 |  |  |  |  |  |  |
| 3C | ywR/ywR;nSyb-Gal80/+;NPF-Gal4/UAS-attp2 | 10% sugar<br>2% yeast | 487 | 66.1232 | 0.39543 | 66 | . | . |  |  |  |
| 3C | ywR/ywR;nSyb-Gal80/+;NPF-Gal4/UAS-NPF-RNAi | 10% sugar<br>2% yeast | 491 | 68.0326 | 0.44922 | 70 | . | . | Log-Rank | 35.5501 | <0.0001 |
| 3C | ywR/ywR;nSyb-Gal80/+;NPF-Gal4/UAS-attp2 | 10% sugar<br>10% yeast | 478 | 56.6736 | 0.41983 | 58 | 58 | 60 |  |  |  |
| 3C | ywR/ywR;nSyb-Gal80/+;NPF-Gal4/UAS-NPF-RNAi | 10% sugar<br>10% yeast | 331 | 60.5619 | 0.40318 | 62 | 60 | 62 | Log-Rank | 41.7776 | <0.0001 |
| Term |  | Estimate | Std Error | Lower 95% | Upper 95% |  |  |  | Proportional hazard test |  |  |
| Diet[2% Yeast] | | -0.6187126 | 0.026854 | -0.671346 | -0.566066 | | | | Interaction | $\chi^2$ | p |
| Genotype[nSybGal80,NPF>attp2] |  | 0.18976 | 0.024195 | 0.1424101 | 0.237272 |  |  |  | Diet | 506.464349 | <0.0001 |
| Genotype[nSybGal80,NPF>attp2]*Diet[2% Yeast] |  | -0.0344553 | 0.024075 | -0.081732 | 0.012657 |  |  |  | Genotype | 61.9840542 | <0.0001 |
|  |  |  |  |  |  |  |  |  | Diet*Genotype | 2.05313435 | 0.1519 |

| Fig. | Genotype | RU486 | Sample size | Mean | Mean s.e. | Median (d) | Lower 95% | Upper 95% | Test | $\chi^2$ | p |
| --- | --- | --- | --- | --- | --- | --- | --- | --- | --- | --- | --- |
| 4A | ywR/ywR;Dilp2GS-Gal4/UAS-NPF-R-RNAi;+/+ | 0 | 476 | 47.4538 | 0.433 | 48 | 48 | 50 |  |  |  |
| 4A | ywR/ywR;Dilp2GS-Gal4/UAS-NPF-R-RNAi;+/+ | 200 | 485 | 59.8763 | 0.394 | 58 | 56 | 58 | Log-Rank | 335.2444 | <0.0001 |
| 4B | ywR/ywR;S106GS-Gal4/UAS-NPF-R-RNAi;+/+ | 0 | 481 | 62.9896 | 0.383 | 64 | 64 | 66 |  |  |  |
| 4B | ywR/ywR;S106GS-Gal4/UAS-NPF-R-RNAi;+/+ | 200 | 479 | 62.5929 | 0.426 | 64 | . | . | Log-Rank | 0.327 | 0.5674 |
| 4C | ywR/ywR;UAS-VDRC-60100/+;nos-VP16-Gal4/+ |  | 476 | 48.6176 | 0.406 | 50 | 48 | 50 |  |  |  |
| 4C | ywR/ywR;UAS-NPF-R-RNAi/+;nos-VP16-Gal4/+ |  | 472 | 46.4746 | 0.37 | 48 | 46 | 48 | Log-Rank | 28.3621 | <0.0001 |
| 4D | ywR/ywR;AKH-Gal4/VDRC-60100;+/+ |  | 504 | 48.2579 | 0.375 | 48 | 48 | 50 |  |  |  |
| 4D | ywR/ywR;AKH-Gal4/UAS-NPF-R-RNAi;+/+ |  | 461 | 50.3601 | 0.443 | 50 | 50 | 52 | Log-Rank | 17.5829 | <0.0001 |

| Fig | Genotype | Treat | Sample size | Mean | Mean s.e. | Median (d) | Lower 95% | Upper 95% | Test | $\chi^2$ | p |
| --- | --- | --- | --- | --- | --- | --- | --- | --- | --- | --- | --- |
| 5D | ywR/ywR;nSyb-Gal80/+;NPF-Gal4/UAS-attp2 | EtOH | 476 | 64.6471 | 0.46397 | 66 | 66 | 68 |  |  |  |
| 5D | ywR/ywR;nSyb-Gal80/+;NPF-Gal4/UAS-NPF-RNAi | EtOH | 472 | 68.1483 | 0.3765 | 70 | 68 | 70 | Log-Rank | 33.4546 | <0.0001 |
| 5D | ywR/ywR;nSyb-Gal80/+;NPF-Gal4/UAS-attp2 | JH | 480 | 43.1542 | 0.722 | 46 | 42 | 46 |  |  |  |
| 5D | ywR/ywR;nSyb-Gal80/+;NPF-Gal4/UAS-NPF-RNAi | JH | 471 | 38.5648 | 0.77904 | 38 | 34 | 42 | Log-Rank | 7.5601 | 0.006 |
| Term |  | Estimate | Std Error | Lower 95% | Upper 95% |  |  |  | Proportional hazard test |  |  |
| Genotype[ nSyb-Gal80,NPF>attp2] | | 0.04804394 | 0.0232176 | 0.0025429 | 0.0935693 | | | | Interaction | $\chi^2$ | p |
| Treat[EtOH] |  | -0.7356221 | 0.0250372 | -0.784693 | -0.686538 |  |  |  | Genotype | 4.28279218 | 0.0385 |
| Genotype[nSyb-Gal80,NPF>attp2]*Treat[EtOH] |  | 0.08638556 | 0.0230709 | 0.041164 | 0.1316154 |  |  |  | Treat | 817.893157 | <0.0001 |
|  |  |  |  |  |  |  |  |  | Treat*Genotype | 14.0078618 | 0.0002 |

| Fig | Genotype | Treat | RU486 | Sample size | Mean | Mean s.e. | Median (d) | Lower 95% | Upper 95% | Test | $\chi^2$ | p |
| --- | --- | --- | --- | --- | --- | --- | --- | --- | --- | --- | --- | --- |
| 5E | ywR/ywR;Dilp2-GS-Gal4/UAS-NPF-R-RNAi;+/+ | EtOH | 0 | 476 | 62.3193 | 0.47642 | 60 | 60 | 62 |  |  |  |
| 5E | ywR/ywR;Dilp2-GS-Gal4/UAS-NPF-R-RNAi;+/+ | EtOH | 200 | 491 | 70.8391 | 0.43851 | 72 | 70 | 72 | Log-Rank | 112.8884 | <0.0001 |
| 5E | ywR/ywR;Dilp2-GS-Gal4/UAS-NPF-R-RNAi;+/+ | JH | 0 | 481 | 57.1892 | 0.4214 | 56 | 56 | 58 |  |  |  |
| 5E | ywR/ywR;Dilp2-GS-Gal4/UAS-NPF-R-RNAi;+/+ | JH | 200 | 482 | 59.2656 | 0.48978 | 60 | . | . | Log-Rank | 15.8203 | <0.0001 |
| Term |  | Estimate | Std Error | Lower 95% | Upper 95% |  |  |  |  | Proportional hazard test |  |  |
| RU[0RU] | | 0.21186727 | 0.0229241 | 0.1669255 | 0.2568012 | | | | | Interaction | $\chi^2$ | p |
| Treat[EtOH] |  | -0.3848803 | 0.0240957 | -0.432142 | -0.337675 |  |  |  |  | RU | 84.736211 | <0.0001 |
| RU[0RU]*Treat[EtOH] |  | 0.0948005 | 0.0228655 | 0.0499747 | 0.1396207 |  |  |  |  | Treat | 252.746757 | <0.0001 |
|  |  |  |  |  |  |  |  |  |  | RU*Treat | 17.1595547 | <0.0001 |

| Fig | Genotype | Treat | Sample size | Mean | Mean s.e. | Median (d) | Lower 95% | Upper 95% | Test | $\chi^2$ | p |
| --- | --- | --- | --- | --- | --- | --- | --- | --- | --- | --- | --- |
| 5G | ywR/ywR;Aug21-Gal4/+;+/+ | EtOH | 470 | 52.3532 | 0.47329 | 54 | 54 | 56 |  |  |  |
| 5G | ywR/ywR;Aug21-Gal4/+;UAS-InR-DN/+ | EtOH | 486 | 60.8313 | 0.41739 | 62 | 62 | 64 | Log-Rank | 245.5575 | <0.0001* |
| 5G | ywR/ywR;Aug21-Gal4/+;+/+ | JH | 348 | 52.6552 | 0.63734 | 55 | 54 | 56 |  |  |  |
| 5G | ywR/ywR;Aug21-Gal4/+;UAS-InR-DN/+ | JH | 471 | 54.6327 | 0.5838 | 58 | 56 | 58 | Log-Rank | 12.8607 | 0.0003 |
| Term |  | Estimate | Std Error | Lower 95% | Upper 95% |  |  |  | Proportional hazard test |  |  |
| Genotype[Aug21>InR-DN] | | -0.2779821 | 0.024431 | -0.325809 | -0.230026 | | | | Interaction | $\chi^2$ | p |
| Treat[EtOH] |  | -0.0731685 | 0.024038 | -0.120225 | -0.025981 |  |  |  | Genotype | 125.984517 | <0.0001 |
| Genotype[Aug21>InR-DN]*Treat[EtOH] |  | -0.1562301 | 0.024159 | -0.203652 | -0.108932 |  |  |  | Treat | 9.21711885 | 0.0024 |
|  |  |  |  |  |  |  |  |  | Genotype*Treat | 42.0093239 | <0.0001 |

**Table S2.**
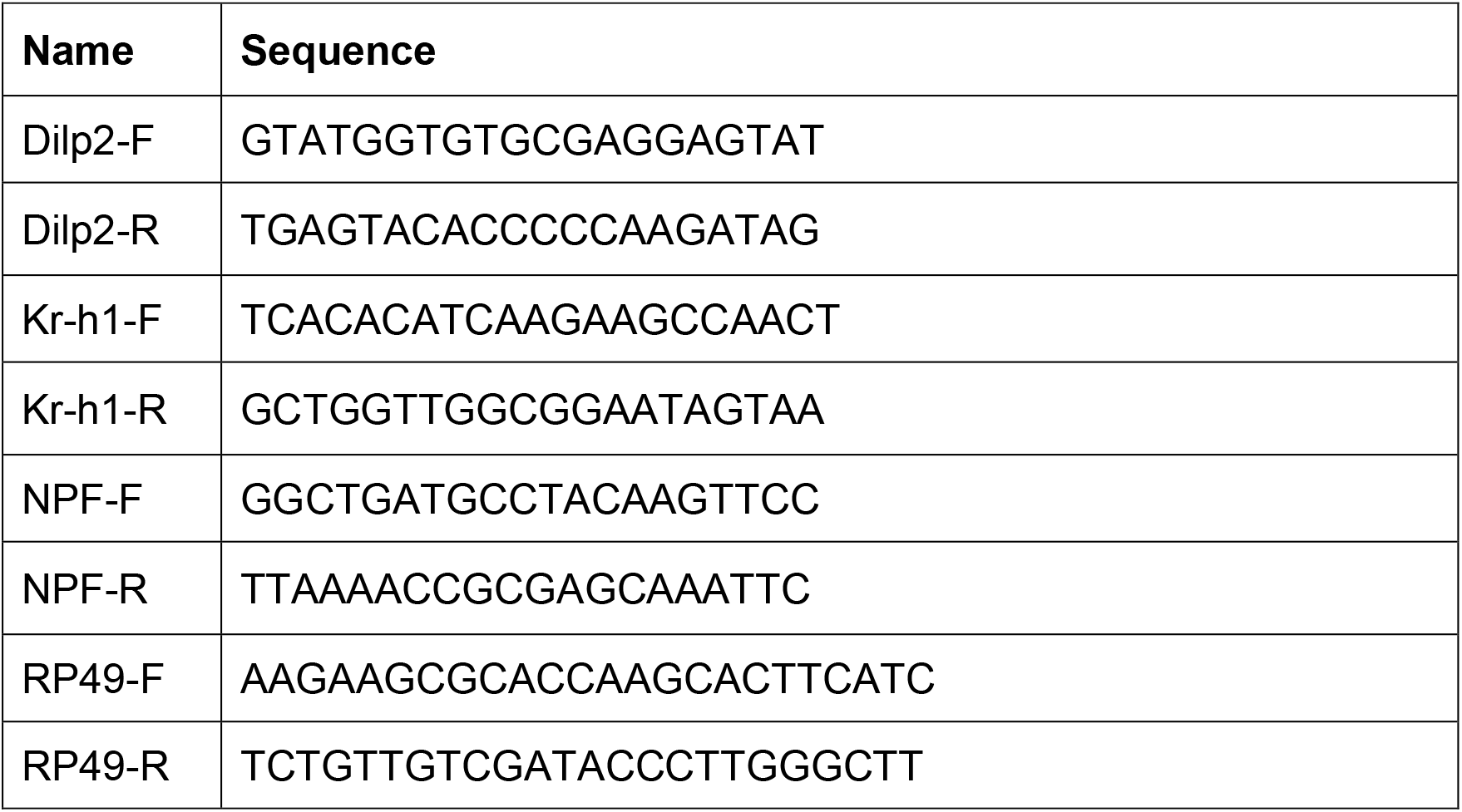
RT-PCR primer.

